# The genetic etiology of longitudinal measures of predicted brain ageing in a population-based sample of mid to late-age males

**DOI:** 10.1101/2021.08.04.455143

**Authors:** Nathan A. Gillespie, Sean N. Hatton, Donald J Hagler, Anders M. Dale, Jeremy A. Elman, Linda K. McEvoy, Lisa T. Eyler, Christine Fennema-Notestine, Mark W. Logue, Ruth E. McKenzie, Olivia K. Puckett, Xin M. Tu, Nathan Whitsel, Hong Xian, Chandra A. Reynolds, Matthew S. Panizzon, Michael J. Lyons, Michael C. Neale, William S. Kremen, Carol Franz

## Abstract

Magnetic resonance imaging data are being used in statistical models to predicted brain ageing (PBA) and as biomarkers for neurodegenerative diseases such as Alzheimer’s Disease. Despite their increasing application, the genetic and environmental etiology of global PBA indices is unknown. Likewise, the degree to which genetic influences in PBA are longitudinally stable and how PBA changes over time are also unknown. We analyzed data from 734 men from the Vietnam Era Twin Study of Aging with repeated MRI assessments between the ages 52 to 72 years. Biometrical genetic analyses ‘twin models’ revealed significant and highly correlated estimates of additive genetic heritability ranging from 59% to 75%. Multivariate longitudinal modelling revealed that covariation between PBA at different timepoints could be explained by a single latent factor with 73% heritability. Our results suggest that genetic influences on PBA are detectable in midlife or earlier, are longitudinally very stable, and are largely explained by common genetic influences.

**Highlights:** We explored the genetic and environmental etiology of MRI-based predicted brain age (PBA) in a longitudinal sample of males starting in midlife. Genetic influences on PBA are detectable in midlife or earlier, are longitudinally very stable, and largely explained by common genetic influences.

## 1. Introduction

Brain magnetic resonance imaging (MRI) data are increasingly used to model predicted brain ageing (PBA). The models assume that MRI of neuroanatomical degeneration reflects poorer brain health and risk of neurodegenerative diseases such as Alzheimer’s Disease (Cole et al., 2019; McEvoy et al., 2009; Wang et al., 2019). The modelling relies on machine learning to estimate associations between MRI data and chronological age in training samples of varying age (Cole and Franke, 2017). These associations are then applied using supervised learning algorithms to estimate PBA or predicted brain age difference (PBAD) (the difference between predicted and chronological age) in independent samples. Broadly, this approach assumes that individual differences in brain aging stem from biological processes influencing lifespan and age-related diseases, which can be explained by genetic and environmental influences (Cole et al., 2019). This assumption has been partly supported by two twin studies reporting the heritability of PBA and PBAD; Cole et al.’s (Cole et al., 2017) cross-sectional analysis of 62 female twins at mean age 62 years and Brouwer et al.’s (Brouwer et al., 2021) longitudinal analysis of 673 twins aged 10 to 23 years. The latter reported PBAD heritabilities up to 79% and longitudinal genetic correlations ranging 0.46 to 0.68 based on grey matter density and cortical thickness. Although this suggests a combination of stable and age-varying genetic processes influencing aging in adolescents and very young adults, we are unaware of any twin studies that have explored the heritability and longitudinal stability of genetic and environmental risks in PBA or PBAD beginning middle-through to later-age when neurodegenerative diseases typically begin to emerge.

Not only can measures of brain ageing predict cognitive decline and morbidity (Cole et al., 2019; Elliott et al., 2019; Franke et al., 2012; Liem et al., 2017), biologically ‘older’ brains are linked to older facial appearance, early cognitive decline including the progression from mild cognitive impairment to dementia, Alzheimer’s Disease, accelerated ageing and shorter lifespans (Cole et al., 2019; de Lange and Cole, 2020; Deary et al., 2009; Elliott et al., 2019; Fjell et al., 2014; Gaser et al., 2013; Lowe et al., 2016; Salthouse, 2010; Vos et al., 2012). Our team has demonstrated associations between negative life events (e.g. family deaths, financial problems, unemployment) and advanced PBA (Hatton et al., 2018a). However, hypotheses regarding heritability and the stability of genetic influences in PBA remain untested.

Twin studies have demonstrated moderate to high heritability in cortical and subcortical volume (Baare et al., 2001; Brouwer et al., 2014; Kremen et al., 2010; Peper et al., 2007; Renteria et al., 2014; Satizabal et al., 2019; Wright et al., 2002), cortical thickness (Kremen et al., 2013a; Kremen et al., 2010; Thompson et al., 2001; Vuoksimaa et al., 2015), cortical surface area (Brouwer et al., 2014; Eyler et al., 2011; Kremen et al., 2013a; Kremen et al., 2010; Vuoksimaa et al., 2015), and diffusion MRI metrics (Elman et al., 2017; Gillespie et al., 2017; Hatton et al., 2018b). Based on these findings including those of Cole et al. (Cole et al., 2017) and Brouwer et al. (Brouwer et al., 2021) we hypothesized that whole brain indicators of PBA based on supervised machine learning combinations of cortical thickness, surface area and subcortical volume metrics in adults should also be heritable. Given the pivot towards early detection of neurodegenerative disease (Albert et al., 2011; Daviglus et al., 2010; Golde et al., 2011; Sperling et al., 2011a; Sperling et al., 2011b), an accurate description of the genetic and environmental etiology of brain aging beginning middle-age is required. Questions that have not been addressed include, how do genetic and environmental risks in PBA develop over time and are they longitudinally stable versus age-specific? Similarly, are PBA genetics in middle-and later-age quantitatively or qualitatively distinct? It is also unclear if genetic and environmental influences, protective or detrimental, accumulate and continue to exert an impact over time. Hypotheses of ageing such as somatic mutation theory predict an accumulation of unrepaired cellular and molecular damage arising from genome instability during a single generation (Kirkwood, 1977, 2005; Morley, 1998). This is consistent with an autoregression model (Boomsma et al., 1989; Boomsma and Molenaar, 1987; Eaves et al., 1986; Guttman, 1954). If changes in brain ageing do stem from an accumulation of age-related genetic and environmental risks, the task, therefore, is to determine how well autoregression explains PBA data. Plausibly, genetic and environmental differences in PBA across time might be better explained by common or independent pathway theories positing time-independent risks (Neale and Cardon, 1992). Our aim, therefore, was to explore the etiology of PBA in a sample of middle-to later-age men with longitudinal MRI assessments. In addition to estimating PBA heritability, we tested competing hypotheses to determine which best explains changes in the genetic and environmental influences in PBA.

## 2. Materials and methods

### 2.1. Subjects

Participants comprise middle-aged male twins who underwent MRI scanning as part of the Vietnam Era Twin Study of Aging (VETSA) (Kremen et al., 2013b). Wave 1 took place between 2001-2007 (Kremen et al., 2006) (mean age=56.1, SD=2.6, range=51.1 to 60.2). Wave 2 occurred approximately 5.5 years later (mean age=61.8, SD=2.6, range=56.0 to 65.9). Wave 3 occurred approximately 5.7 years later (mean age=67.5, SD=2.6, range=61.4 to 71.7). All participants were concordant for US military service at some time between 1965-1975, but nearly 80% reported no combat experience. The sample is 88.3% non-Hispanic white, 5.3% African-American, 3.4% Hispanic, and 3.0% “other” participants. Based on data from the US National Center for Health Statistics, the sample is very similar to American men in their age range with respect to health and lifestyle characteristics (Schoeneborn and Heyman, 2009). Written informed consent was obtained from all participants.

### 2.2. Ethics statement

The University of California, San Diego, Human Research Protection Program Institutional Review Board approved the proposal to collect these data (Project #150572, 150572, 150537, 140361, 071446, 031639, 151333). Data are publicly available through requests at the VETSA website (http://www.vetsatwins.org).

### 2.3. Magnetic resonance imaging acquisition and analysis

At Wave 1, MR images were acquired on Siemens 1.5 Tesla scanners (N=260 at University of California, San Diego (Siemens-Symphony); N=226 at Massachusetts General Hospital: MGH; Siemens-Avanto). Sagittal T1-weighted magnetization-prepared rapid gradient echo (MPRAGE) sequences were employed with TI=1000 ms, TE=3.31 ms, TR=2730 ms, flip angle=7 degrees, slice thickness=1.33 mm, voxel size 1.3×1.0×1.3 mm. Raw DICOM MRI scans (including two T1-weighted volumes per case) were downloaded to the MGH site.

At Wave 2, T1-weighted images providing high anatomical detail were acquired on 3T scanners at University of California, San Diego (UCSD) and Massachusetts General Hospital. At UCSD, images were acquired on a GE 3T Discovery 750scanner (GE Healthcare, Waukesha, WI, USA) with an eight-channel-phased array head coil. The imaging protocol included a sagittal 3D fast-spoiled gradient echo T1-weighted image (echo time=3.164 msec, repetition time=8.084 msec, inversion time=600 msec, flip angle=8 degrees, pixel bandwidth=244.141, field of view=25.6 cm, frequency=256, phase=192, slices=172, and slice thickness=1.2 mm) At Massachusetts General Hospital, images were acquired with a Siemens Tim Trio, (Siemens USA, Washington, D.C.) with a 32-channel head coil. The imaging protocol included a 3D MPRAGE T1-weighted image (echotime 4.33 msec, repetition time 2170 msec, inversion time 1100 msec,flip angle 7 degrees, pixel bandwidth 140, field of view 25.6 cm, frequency 256, phase 256, slices 160, and slice thickness 1.2 mm).

At Wave 3, all participants were scanned at UCSD. The acquisition protocol was identical to the protocol used at UCSD during Wave 2.

As described in prior work (Eyler et al., 2012), raw image files from Wave 1, 2 and 3 were processed using the same in-house pipeline written in MATLAB and C by the UCSD Center for Multimodal Imaging and Genetics. Data were qualitatively assessed and images with severe scanner artifacts or excessive head motion were either rescanned where possible or excluded from the analysis (approximately 3%). T1-weighted structural images were corrected for gradient distortions (Jovicich et al., 2006) and B1 field inhomogeneity (Sled et al., 1998). Subcortical segmentation and surface-based cortical parcellation were performed using FreeSurfer, Version 5.3 (Fischl, 2012). Inaccuracies in cortical surfaces were manually corrected by trained neuroimaging analysts. All images required some form of manual editing to ensure the correct classification of the pial and white matter surfaces, with particular attention given to the orbitofrontal cortex, the temporal lobes, meninges, and transverse and superior sagittal sinuses. Problematic segmentations/parcellations were reviewed by consensus with four neuroimaging analysts.

### 2.4. Predicted brain age & predicted brain age (PBA) difference endophenotypes

Primary analyses focused on PBA. Discussed in detail elsewhere (Hatton et al., 2018a), PBA was estimated using the Brain-Age Regression Analysis and Computation Utility software BARACUS v0.9.4 (2017; Liem et al.). BARACUS uses linear support vector regression models to predict brain age derived from each individual’s FreeSurfer statistics. Specifically, vertex-wise cortical metrics were derived from the fsaverage4 standard space for cortical thickness (n=5124 vertices) and surface area (n=5124 vertices), and subcortical segmentation metrics were derived from the aseg.stat file for subcortical volume (n=66 regions of interest). We used the BIDS-mode docker on Ubuntu 16.04 using the default database (Liem2016_OCI_norm), which is trained on 1166 subjects with no objective cognitive impairment (566 female/600 male, mean age 59.1 years, SD 15.2, range 20-80 years).

In secondary analyses, we considered predicted brain age difference (PBAD) scores, which were calculated by subtracting PBA (referred to as “stacked-anatomy” brain age in BARACUS) from the chronological age. A negative PBAD is indicative of brain age estimated to be older than one’s chronological age. We note that while supervised machine learning algorithms such as BARACUS can detect informative multivariate patterns, the relative contributions of individual regions are not tested. Therefore, no inferences are made regarding particular regions driving PBA/PBAD.

As noted in Section 2.1., there was considerable variation in chronological age at each wave and overlap in age ranges between the three assessments. Given the variation and overlap, longitudinal analysis of these wave-based data would therefore preclude any meaningful understanding of age-related changes. Ignoring irregular spacing between time intervals in longitudinal modeling can lead to biased parameter estimates (Estrada and Ferrer, 2019). Rather than employing definition variables to account for individual differences in age at assessment and irregular timer intervals (Mehta and Neale, 2005), our solution was to recode each subject’s score according to their chronological age at assessment. Thus, for example, if two subjects ‘a’ and ‘b’ were both aged 60 at VETSA 1 and 2 respectively, each would be assigned a PBA score for age 60. Since each subject contributed a maximum of three data points between ages 51 to 72, this creates missing data for which Full Information Maximum Likelihood is well suited to handling. However, to reduce sparse data while maintaining computational efficiency, our ‘age-anchored’ PBA and PBAD scores were re-coded to one of four age intervals according to each individual’s age at assessment: 51 to 55 years; 56 to 60 years; 61 to 65 years; and 66 to 72 years.

There were 260, 251, and 126 subjects with PBA scores at one, two and three age intervals respectively. Since there were only 3 VETSA assessments, no subjects had data from all four age intervals. Five participants were ascertained twice in the same five-year age interval. Only their first observation was included. Prior to twin modelling all PBA and PBAD scores were residualized for the effects of scanner (i.e., 1.5T vs 3T) and ethnicity using the umx_residualize function in the umx software package (Bates et al., 2019), and given the range in birth year (1943 to 1955), residuals were also adjusted for cohort effects.

### 2.5. Statistical analyses

The OpenMx_2.9.9.1_ software package (Boker et al., 2011) in R_3.4.1_ (R Development Core Team, 2018) was used to estimate correlations between the PBA scores and to fit univariate and multivariate genetic twin models (Neale and Cardon, 1992). OpenMx software coding used for the multivariate analyses is included in the Supplement. Given the numbers of incomplete twin pairs (see Supplement Table S1), methods such as Weighted Least Squares would result in significant listwise deletion thereby altering the accuracy of the PBA and PBAD means and variances. Fortunately, the raw data Full Information Maximum Likelihood (FIML) option in OpenMx_2.9.9.1_ (Boker et al., 2011) has the advantage of not only being robust to violations of non-normality but also enables analysis of missing or incomplete data as well as the direct estimation of covariate effects. More accurate means and variance improve the estimation of the variances and covariance structure used to test our competing hypotheses.

Figure 1

**Figure 1.**
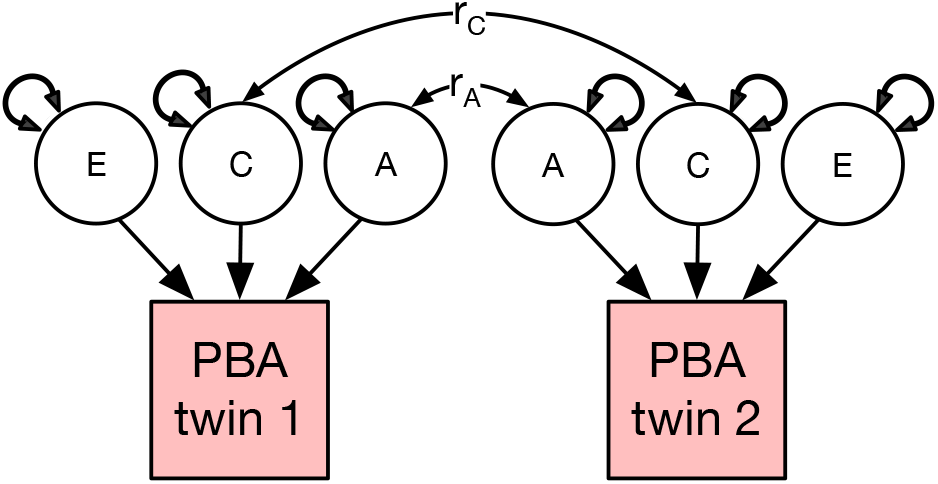
Univariate variance decomposition to estimate the relative contribution of genetic & environmental influences in predicted brain ageing (PBA). A = additive genetic, C = common / shared environmental, and E = unshared environmental influences, r_C_ = correlation of 1 for MZ and DZ twin pairs, r_A_ = correlations 1 for MZ and 0.5 for DZ twin pairs.

### 2.6. Univariate analyses

In univariate analyses, the total variation in each PBA score was decomposed into additive (A) heritability, shared or common environmental (C), and non-shared or unique (E) environmental variance components (see Figure 1). This approach is referred to as the ‘ACE’ variance component model. The decomposition is achieved by exploiting the expected genetic and environmental correlations between MZ and DZ twin pairs. MZ twin pairs are genetically identical, whereas DZ twin pairs share, on average, half of their genes. Therefore, the MZ and DZ twin pair correlations for the additive genetic effects are fixed to r_A_=1.0 and r_A_=0.5 respectively. The modelling assumes that shared environmental effects (C) are equal in MZ and DZ twin pairs (r_C_=1.0), while non-shared environmental effects (E) are by definition uncorrelated and include measurement error.

### 2.7. Multivariate analyses to test competing theories

This univariate method is easily extended to the multivariate case to estimate the size and significance of genetic and environmental influences within and between PBA over time. In order to have a reference for contrasting and choosing the best fitting model, we first fitted a multivariate ACE ‘correlated factors’ (Figure 2a). This is a saturated model that reproduces perfectly all mean and variance-covariance information for the observed PBA variables.

**Figure 2.**
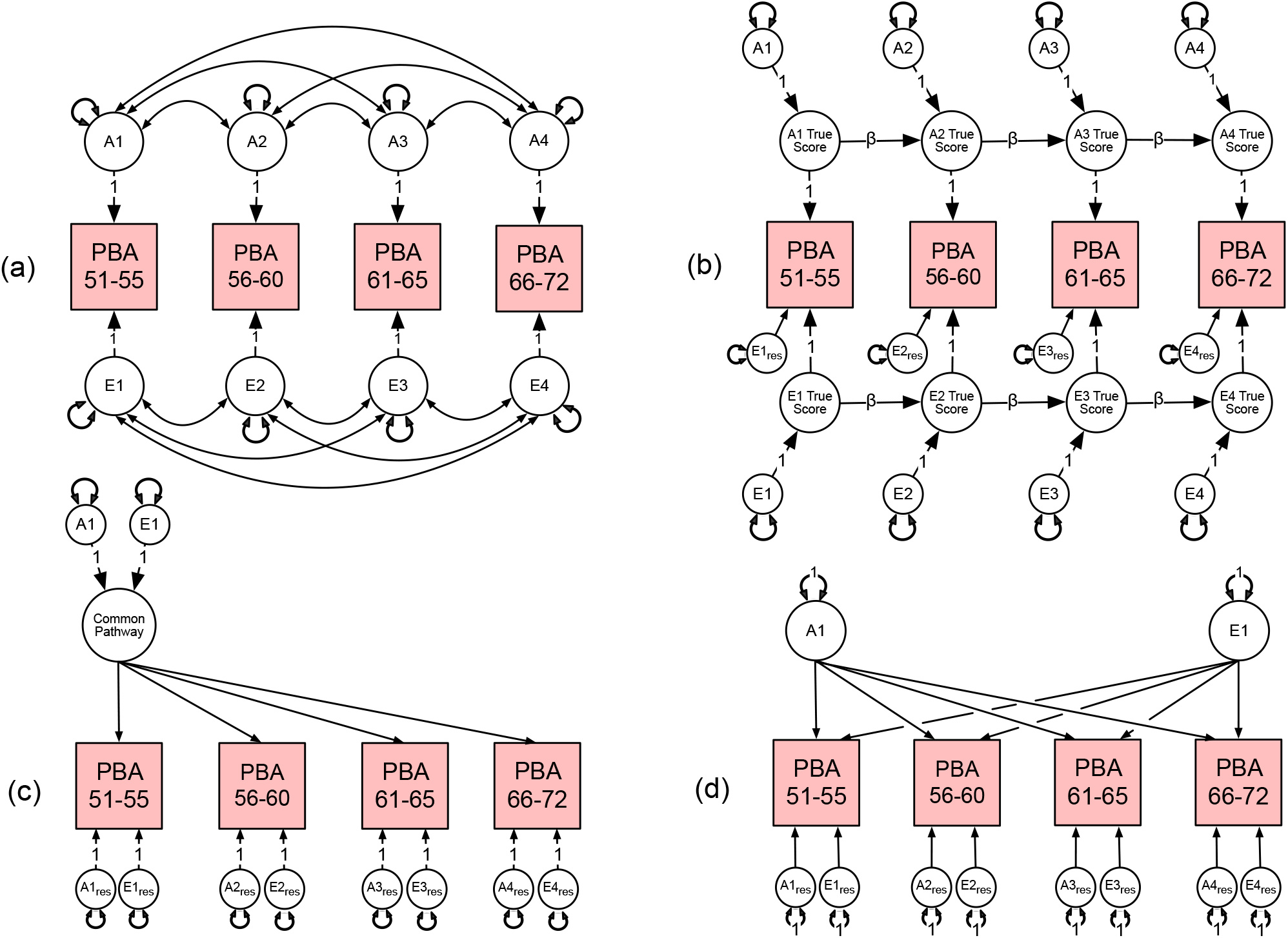
Multivariate correlated factors (a) and competing hypothetical models to explain the sources of variance-covariance between the predicted brain age (PBA) scores. These include (b) the auto-regression, (c) common pathway model, and (d) independent pathway models. For brevity, only additive genetic (A) & non-shared environmental (E) factors are shown.

Figure 2

The ACE correlated factors model makes no theoretical prediction regarding how genes and environments change over time. In contrast, the autoregression model (Figure 2b) predicts that time-specific random genetic or environmental effects may persist over time (autoregressive effects) (Eaves et al., 1986). As described elsewhere (Boomsma et al., 1989; Boomsma and Molenaar, 1987; Eaves et al., 1986), autoregression assumes that a trait measured at time *t* is partly a function of the same trait measured at a prior timepoint *t-1*. New variation at each assessment reflects time-specific genetic or environmental influences. Such autoregression may occur between phenotypes, or between the latent genetic or environmental true scores. The variance in the genetic and environmental true scores on each occasion is a function of (i) new random genetic and environmental effects or innovations arising at each time point and (ii) the causal contribution, via the beta coefficient, from the true scores expressed at preceding times. All cross-temporal correlations within subjects arise when the contribution of innovations are more or less persistent over time. These contributions may, under some circumstances accumulate, potentially giving rise to developmental increases in genetic or environmental variance and increased correlations between adjacent measures. Depending on the magnitude of an innovation and its relative persistence, the observed variances and cross-temporal covariances may also increase during development towards a stable asymptotic value. Another feature of the autoregressive model is that cross-temporal correlations tend to decay as a function of increasing time differential. Our modeling also included occasion-specific residual variance including measurement error not captured by the autoregression process. See Eaves et al. (Eaves et al., 1986) for graphical examples of an application to longitudinal cognitive data.

In contrast, the common pathway (CP) model (Figure 2c) predicts a covariance structure between all four PBA scores explained by a common liability decomposed into A, C, and E influences. As with factor analysis, the CP is ‘indicated’ by the strength of the factor loadings to each observed PBA score. Residual variances or risks unique to each PBA score are further decomposed into variable specific genetic and environmental residuals. To further explore the multivariate space, we fitted a model with two common factors.

Finally, the independent pathway model (Figure 2d) predicts that latent genetic and environmental risk factors separately generate covariance between the PBA scores. Again, variances unique to each PBA score not captured by these independent pathways are decomposed into variable specific genetic and environmental residuals.

### 2.8. Model fit

The best-fitting model was determined using a using a likelihood ratio test and the Akaike’s Information Criterion (AIC) (27). For each best-fitting univariate and multivariate model, the parameters were then successively fixed to zero and their significance determined using a likelihood ratio chi-square test. We have argued elsewhere that the advantage of AIC is its deep theoretical connections to cross-validation (Kirkpatrick et al., 2015). Specifically, in large samples, the AIC is expected to select that model in the candidate set which minimizes the error of prediction in new samples of the same size from the population (where the error is based on a log-likelihood function) (Kirkpatrick et al., 2015). Specifically, the AIC is expected to minimize the Kullback–Leibler (KL) divergence from full reality at the given sample size. A sensible objective of model selection is to choose the model that has the smallest KL divergence from full reality. The full reality, of course, is not known, and may not even be knowable. Indeed, a complete description of full reality would be infinitely long. However, if we accept the possibility that no statistical model can completely describe reality, then the premise of there being a ‘true model’ that generated the data becomes rather dubious. In summary, because full reality may be unknowable, we do not presume that the true model is knowable from our data and consequently, chose our fit index based on this philosophy. Rather than proposing to identify the true model, the AIC selects the best-approximating model based on an optimal balance of parsimony and model fit.

## 3. Results

The numbers of complete and incomplete twin pairs by zygosity are shown in Supplementary Table S1. Descriptive statistics for each PBA score before and after residualization of the means and variances are shown in Supplementary Table S2.

### 3.1. Strength of association

As shown in Table 1, the phenotypic correlations between the age-anchored PBA scores at each age interval were high and ranged from 0.67 to 0.76.

**Table 1.** Pairwise polyserial phenotypic correlations for the predicted brain age (PBA) scores. Polyserial correlations represent the associations between the underlying liability rather than observed phenotypic distributions (Pearson, 1900; Pearson and Pearson, 1922).

|  | 1. | 2. | 3. | 4. |
| --- | --- | --- | --- | --- |
| 1. PBA 51 to 55 | 1 |  |  |  |
| 2. PBA 56 to 60 | 0.67 | 1 |  |  |
| 3. PBA 61 to 65 | 0.76 | 0.74 | 1 |  |
| 4. PBA 66 to 72 | 0.67 | 0.72 | 0.75 | 1 |

### 3.2. Twin pair correlations

Table 2 shows the twin pair correlations by zygosity for PBA at each age interval. If familial aggregation was entirely attributable to shared family environments, then monozygotic (MZ) and dizygotic (DZ) twin pair correlations would be statistically equal. In contrast, if familial aggregation was entirely attributable to shared additive (or non-additive) genetic factors, then DZ correlations would be ½ (or less) the size of the MZ twin pair correlations. Here, DZ twin pair correlations ranged from r_dz_=0.1 to r_dz_=0.6 and were approximately 1/3 the size of the MZ twin pair correlations. This is consistent with familial aggregation attributable to genetic risks.

**Table 2.** Monozygotic & dizygotic twin pair polyserial correlations (corrMZ & CorrDZ) along with standardized variance components and 95% confidence intervals components for the best-fitting additive genetic (A) & non-shared environment (E) univariate models.

|  | corrMZ | (95%CI) | CorrDZ | (95%CI) | A | (95%CI) | E | (95%CI) |
| --- | --- | --- | --- | --- | --- | --- | --- | --- |
| PBA 51 to 55 | 0.68 | (0.53-0.78) | 0.13 | (-0.19-0.42) | 0.64 | (0.54-0.69) | 0.36 | (0.31-0.46) |
| PBA 56 to 60 | 0.70 | (0.59-0.78) | 0.29 | (0.11-0.46) | 0.71 | (0.61-0.79) | 0.29 | (0.21-0.39) |
| PBA 61 to 65 | 0.60 | (0.49-0.70) | 0.20 | (0.01-0.38) | 0.58 | (0.45-0.68) | 0.42 | (0.32-0.55) |
| PBA 66 to 72 | 0.66 | (0.55-0.75) | 0.18 | (-0.07-0.58) | 0.61 | (0.47-0.71) | 0.39 | (0.29-0.53) |

### 3.3. Univariate analyses

Full univariate model fitting results for each PBA score are shown Table S3. At each interval, the ‘AE’ model with no common or shared environmental effects provided the best fit to the data. As summarized in Table 2, familial aggregation in each PBA score could be entirely explained by additive genetic influences (A) ranging from 59% to 75%. All remaining variation was explained by non-shared environmental influences including measurement error (E).

### 3.4. Multivariate analyses

Modelling fit results for the autoregression, 1-and 2-factor common pathway and the independent pathway models including comparisons with the reference ‘ACE’ correlated factors model are shown in Table 3. Both the autoregression and independent pathway models fitted the data poorly as judged by the significant change in their likelihood chi-squared ratios (Δ-2LL) and their higher AIC values. In contrast, the changes in the likelihood for the 2-factor and 1-factor common pathway models were not significant. Here, the later, more parsimonious model provided a better comparative fit as judged by the lower AIC. In subsequent modeling based on this best fitting 1-factor common pathway model (see Supplementary Table S4), both the ‘CE’ and ‘E’ sub-models deteriorated significantly when compared to the saturated ACE whereas the ‘AE’ model yielded a non-significant likelihood ratio chi-square difference as well as the lowest AIC.

**Table 3.** Predicted brain age (PBA) multivariate model fitting results & comparisons between the saturated ACE correlated factors model (reference), autoregression, 1-& 2-factor common pathways, & the independent pathways sub-models. Best fitting model bolded.

| | ep | -2LL | df | Δ-2LL | Δdf | p | AIC |
| --- | --- | --- | --- | --- | --- | --- | --- |
| ACE correlated factors (reference) | 34 | 8117.21 | 1405 | - | - | - | 8185.21 |
| Autoregression | 18 | 8148.24 | 1421 | 31.03 | 16 | 0.013 | 8184.24 |
| 2-factor common pathway | 32 | 8123.46 | 1409 | 6.25 | 4 | 0.181 | 8187.46 |
| **1-factor common pathway** | **23** | **8136.07** | **1417** | **18.86** | **12** | **0.092** | **8182.07** |
| Independent pathways | 28 | 8140.52 | 1411 | 23.31 | 6 | 0.001 | 8196.52 |

Note: ep = number of estimated parameters, -2LL = -2 x log-likelihood, Δ-2LL = change in -2 x log-likelihood, $\Delta$ df = change in degrees of freedom, AIC = Akaike Information Criteria. The ACE correlated factors is useful for ensuring that the observed variance-covariance structure is positive definitive and for evaluating any potential model misspecification (Neale et al., 2006; Verhulst et al., 2019). Under the correlated factors we observed negative estimates for the shared environmental (C) variance components. Although this could reflect either stochastic variation or genetic dominance for which this sample is underpowered to detect (Martin et al., 1978), all 'C' influences could be dropped from model without any significant deterioration in fit such that additive genetic risks explained all sources of familial aggregation in the best-fitting correlated factors.

**Table 3.** Additive genetic (below diagonal) & non-shared environmental correlations based on the best fitting ‘AE’ 1-factor common pathway model.

|  | 1. | 2. | 3. | 4. |
| --- | --- | --- | --- | --- |
| 1. PBA 51 to 55 | 1 | 0.49 | 0.47 | 0.45 |
| 2. PBA 56 to 60 | 0.82 | 1 | 0.60 | 0.58 |
| 3. PBA 61 to 65 | 0.92 | 0.87 | 1 | 0.55 |
| 4. PBA 66 to 72 | 0.83 | 0.78 | 0.88 | 1 |

Figure 3 shows this best-fitting multivariate ‘AE’ single factor or common pathway model. A single common factor, with 73% additive genetic and 27% non-shared environmental variance, best explained the covariance between the PBA scores. In this model, the total genetic variances (common & residual influences) in PBA at ages 51-55, 56-60, 61-65 and 66-72 were 59%, 73%, 59% and 67% respectively. For PBA at ages 61-65, the residual genetic variance was non-significant, indicating that genetic variance here is entirely captured by the common factor.

**Figure 3.**
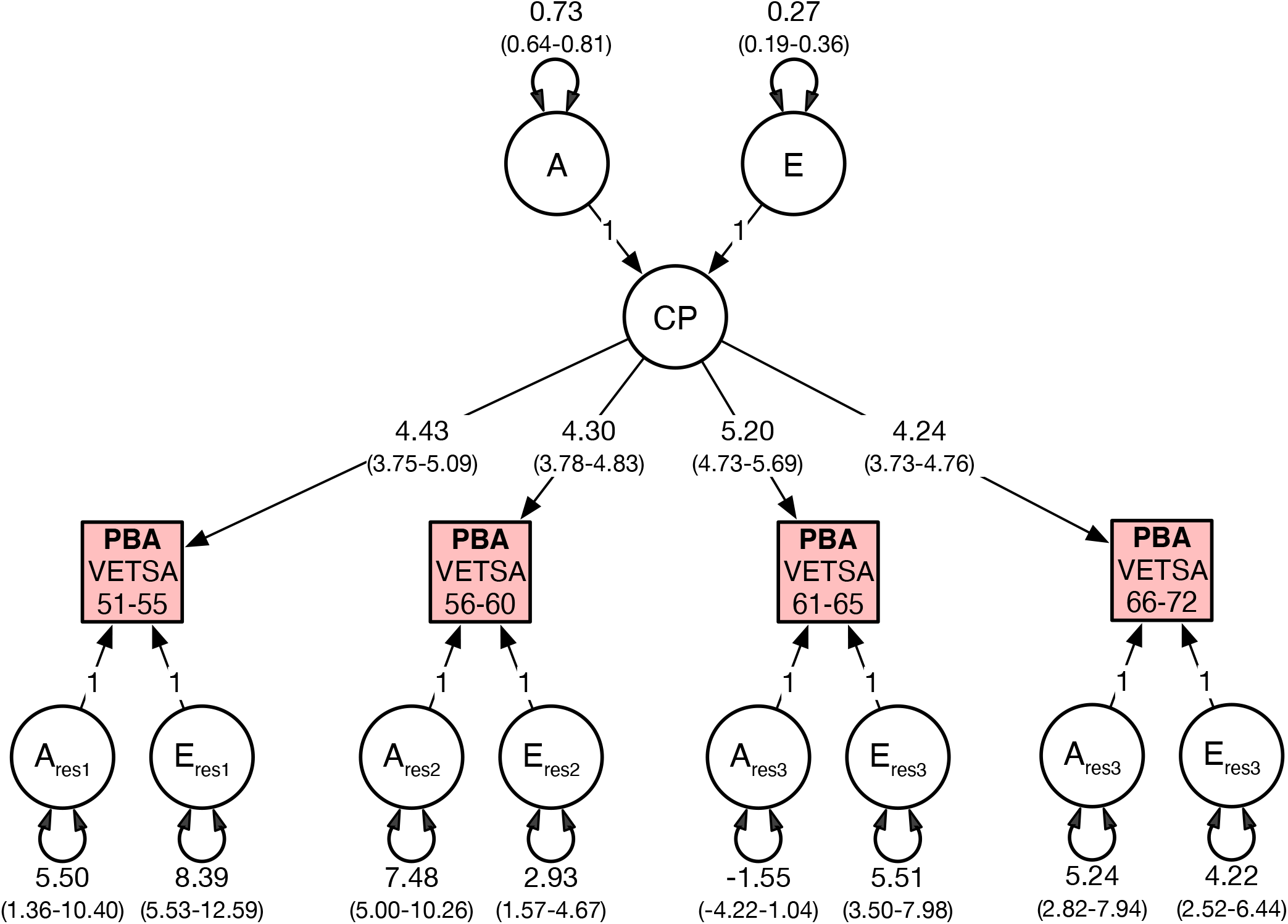
Best fitting common pathway (CP) multivariate model for predicted brain age (PBA) comprising additive genetic (A) and non-shared environment (E) variance components. CP variance components are standardized. All parameters included 95% confidence intervals.

Figure 3

Genetic correlations between the four PBA scores were high and ranged from 0.68 to 0.89 (Table 3) indicating that the same genes are largely influencing PBA across time. In contrast, the environmental correlations were moderate to high, ranging from 0.45 to 0.60 (Table 3) suggesting that large proportions of the environmental influences are unique to each age interval.

We then applied the same univariate and multivariate modelling pipeline to the PBAD scores. Results are shown in Supplementary tables S5-S9. The pairwise phenotypic correlations for the PBAD scores (Supplementary Table S5) were very close to the PBA phenotypic correlations in Table 1. Likewise, the proportion of additive genetic variance in the best fitting univariate PBAD models (Supplementary Table S6) was very similar to the univariate PBA variance components in Table 2. The best-fitting multivariate model was again the single factor common pathway multivariate model (Supplementary Table S7) from which all shared or common environmental ‘C’ effects could be removed without any significant deterioration in the model fit (Supplementary Table S8). Finally, not only were the patterns of additive genetic and non-shared environmental factor correlations in the best fitting 1-factor common pathway ‘AE’ model (Supplementary Table S9) for PBAD nearly identical to those in Table 3, the heritability of the common pathway was again 74% (Supplementary Figure S1).

## 4. Discussion

Individual differences in MRI-based estimates of PBA and PBAD appear highly heritable, with genetic influences accounting for approximately three-quarters of the overall variance. Genetic influences in PBA are also highly correlated across time and are best explained by a common set of risk factors observable in midlife. Consequently, efforts to identify common molecular variants in PBA (Smith et al., 2020) may not require age-stratified samples. Our findings are also consistent with the hypothesis that common genetic risks explain most of the individual differences in brain ageing beginning in midlife and onwards.

Regarding environments, we found that PBA could not be explained by any shared or between-group environmental influences. Instead, all environments were entirely random and unshared and only moderately correlated across time. Thus, in terms of individual differences in brain ageing, environments shared between family members, e.g. similar household incomes and SES (Davies et al., 2015; van der Loos et al., 2013), are of less importance than environments that are unique to individuals, e.g. diet, drug use or allostatic stressors such as negative life events (Hatton et al., 2018a). This finding may have implications concerning the efficacy of community-based versus individually-targeted efforts to slow rates of brain ageing.

Our hypothesis of accumulative environmental and molecular risks predicted by somatic mutation theories that should be captured by autoregression modelling was not supported. Instead, our data were more consistent with what is perhaps a counterintuitive explanation. To the extent that any unrepaired damage is linked to genetic variation in our global indices of PBA, our modelling provided little support for autoregression features or accumulation of age-related or age-specific genetic risks over time. Likewise, we found no evidence to support the hypothesis that age-specific environmental risks are accumulative.

Instead, our best-fitting model suggests that brain ageing is best explained by stable genetic and environmental influences acting via a highly heritable common pathway accounting for most of the individual differences over a 21-year period. Our modelling makes no prediction regarding the number of genes likely involved in brain ageing. Given recent genome wide association scan (GWAS) findings based on multiple brain ageing indices (Smith et al., 2020), including a GWAS of lifespan (Timmers et al., 2019), we speculate that ageing processes are highly polygenic. Our statistically derived common pathway should not be interpreted to represent any identifiable biological structure(s) governing this supervised learning index of ageing. It is, however, consistent with Kirkwood’s theory of a centrally regulated process of ageing, which under selection, has evolved to optimize the “allocation of metabolic resources across core processes like growth, reproduction, and maintenance” (Kirkwood, 2005). Kirkwood et al. also argued that ‘network’ theories of ageing used to describe multiple processes (Kirkwood, 1977, 2005) ought to distinguish upstream mechanisms that set ageing in motion from downstream mechanisms that affect ageing at the cellular level toward the end of life (Kowald and Kirkwood, 1996). The high genetic correlation of r_g_=0.72 between ages 51 to 55 and 66 to 72 suggests that the broad genetic risks underpinning any putative ‘upstream’ and ‘downstream’ processes are mostly shared in common.

### 4.1. Limitations

Our results should be interpreted in the context of four potential limitations.

First, our hypothesis testing was not exhaustive. If PBA is related to rates of cellular or molecular ageing (Kirkwood, 2005), plausibly, genetic and environmental influences could unfold over time, and be better explained by growth processes (Duncan and Duncan, 1991; Duncan et al., 1994; McArdle, 1986; McArdle and Epstein, 1987; Nesselroade and Baltes, 1974). Although each twin pair was assessed on the same scanner on each measurement occasion, MRI data were collected on different scanners (i.e., 1.5T at VETSA 1 vs 3T at VETSA 2 and 3) resulting in likely measurement non-invariance across assessments. Consequently, data were residualized for these and other covariate effects. This resulted in the loss of interpretable mean and variance information necessary for latent growth curve modeling.

Second, data were limited to ages 51 to 72. Plausibly, genetic and environmental autoregression processes occurred before the first assessment (Elliott et al., 2019). There may also exist sub-groups of individuals for whom different autoregressions or hybrid auto-regression plus common factor models provide a better explanation of change. These theoretical alternatives are not within the scope of the current analyses and data.

Third, the age distribution of VETSA spans a decade, but the interval between assessments was less than this, so results should be interpreted as the average change of individuals with this age range. Nevertheless, we repeated our analyses using wave-based data whereby the assessment occasion is treated as a different time point while modeling age at assessment as a covariate. We also reduce the

Finally, our results may not generalize to women or ethnic minorities. We know of no other genetically informative twin studies with comparable and longitudinal MRI data. The uniqueness and size of our sample is its key strength.

### 4.2. Conclusions

This is the first study to explore the genetic and environmental influences on PBA in a longitudinal sample. We assessed males age 51 to 72 years and report three major findings. First, measures of PBA were highly correlated across time. Second, the heritability estimates based on univariate twin analyses ranged from 59% to 74%. Finally, there was no evidence that PBA could be explained by an accumulation of age-specific genetic or environmental risks. Instead, genetic influences at each age interval were highly correlated and captured by a single, common factor with a heritability of 73%. Future analyses should explore the sources of genetic and environmental covariation between brain ageing and other complex behaviors related to cognitive decline.

## Acknowledgements

We would also like to acknowledge the continued cooperation and participation of the members of the VET Registry and their families. The U.S. Department of Veterans Affairs, Department of Defense, National Personnel Records Center, National Archives and Records Administration, Internal Revenue Service, National Opinion Research Center, National Research Council, National Academy of Sciences, and the Institute for Survey Research, Temple University provided invaluable assistance in the conduct of the VET Registry. The Cooperative Studies Program of the U.S. Department of Veterans Affairs provided financial support for development and maintenance of the Vietnam Era Twin Registry.

