## Supplementary information for "The genetic etiology of longitudinal measures of predicted brain ageing in a population-based sample of mid to late-age males"

Figure S1. Best fitting 1-factor common pathway (CP) multivariate model for predicted brain age difference (PBAD) comprising additive genetic (A) & non-shared environment (E) variance components. CP variance components are standardized. All parameters included 95% confidence intervals.

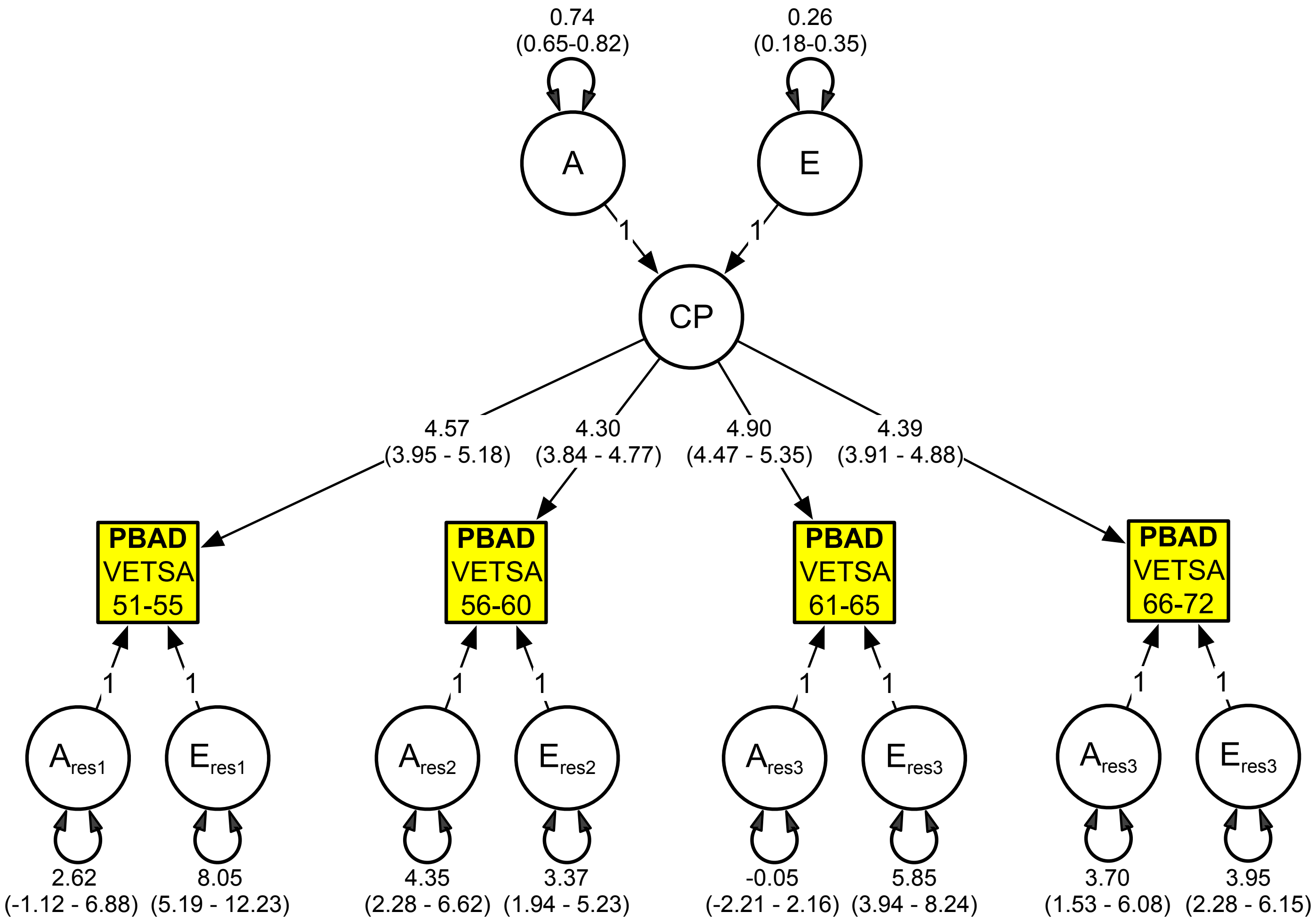

Table S1. Numbers of complete and incomplete monozygotic (MZ) & dizygotic (DZ) twin pairs with predicted brain age (PBA) scores.

| | PBA 51 to 55 | | PBA 56 to 60 | | PBA 61 to 65 | | PBA 66 to 72 | |
|---|---|---|---|---|---|---|---|---|
| | Complete | Singletons | Complete | Singletons | Complete | Singletons | Complete | Singletons |
| MZ | 60 | 20 | 105 | 40 | 98 | 50 | 78 | 51 |
| DZ | 40 | 8 | 82 | 30 | 67 | 37 | 52 | 39 |

Table S2. Phenotypic descriptions for the raw & residualized (adjusted for the effects of scanner, ethnicity & birth year/cohort) predicted brain age (PBA) scores.

| | N | mean | SD | min | max |
|---|---|---|---|---|---|
| PBA 51 to 55 raw | 228 | 63.07 | 6.10 | 45.63 | 74.13 |
| PBA 56 to 60 raw | 442 | 64.80 | 5.19 | 35.22 | 74.86 |
| PBA 61 to 65 raw | 417 | 64.45 | 5.79 | 42.62 | 74.38 |
| PBA 66 to 72 raw | 348 | 66.55 | 5.28 | 42.56 | 75.54 |
| PBA 51 to 55 residualized | 228 | -0.12 | 5.97 | -17.31 | 9.85 |
| PBA 56 to 60 residualized | 442 | 0.10 | 4.91 | -25.06 | 9.71 |
| PBA 61 to 65 residualized | 417 | 0.01 | 5.60 | -21.37 | 9.94 |
| PBA 66 to 72 residualized | 348 | 0.00 | 5.25 | -24.15 | 9.06 |

Table S3. Predicted brain age (PBA) univariate model fitting results.

| PBA 51 to 55 | ep | -2LL | df | Δ-2LL | Δdf | p | AIC |
|---|---|---|---|---|---|---|---|
| ACE | 4 | 1409.34 | 224 | | | | 1417.34 |
| AE | 3 | 1410.68 | 225 | 1.34 | 1 | 0.247 | 1416.68 |
| CE | 3 | 1416.33 | 225 | 6.99 | 1 | 0.008 | 1422.33 |
| E | 2 | 1505.98 | 226 | 96.64 | 2 | <0.001 | 1509.98 |
| PBA 56 to 60 | ep | -2LL | df | Δ-2LL | Δdf | p | AIC |
| ACE | 4 | 2594.15 | 440 | | | | 2602.15 |
| AE | 3 | 2594.47 | 441 | 0.32 | 1 | 0.574 | 2600.47 |
| CE | 3 | 2614.55 | 441 | 20.40 | 1 | <0.001 | 2620.55 |
| E | 2 | 2671.28 | 442 | 77.13 | 2 | <0.001 | 2675.28 |
| PBA 61 to 65 | ep | -2LL | df | Δ-2LL | Δdf | p | AIC |
| ACE | 4 | 2565.84 | 413 | | | | 2573.84 |
| AE | 3 | 2566.84 | 414 | 1.00 | 1 | 0.317 | 2572.84 |
| CE | 3 | 2578.96 | 414 | 13.12 | 1 | <0.001 | 2584.96 |
| E | 2 | 2618.51 | 415 | 52.67 | 2 | <0.001 | 2622.51 |
| PBA 66 to 72 | ep | -2LL | df | Δ-2LL | Δdf | p | AIC |
| ACE | 4 | 2104.14 | 346 | | | | 2112.14 |
| AE | 3 | 2104.16 | 347 | 0.02 | 1 | 0.892 | 2110.16 |
| CE | 3 | 2109.15 | 347 | 5.01 | 1 | 0.025 | 2115.15 |
| E | 2 | 2153.14 | 348 | 48.99 | 2 | <0.001 | 2157.14 |

Note: A = additive genetic, C = common or shared environment, E = non-shared environment, ep = number of estimated parameters, -2LL = -2 x log-likelihood, Δ-2LL = change in -2 x log-likelihood, Δdf = change in degrees of freedom, AIC = Akaike Information Criteria

Table S4. Predicted brain age (PBA) multivariate model fitting results & comparisons between the best fitting ACE 1-factor common pathway model and the AE, CE & E sub-models.

| | ep | -2LL | df | Δ-2LL | Δdf | p | AIC |
|---|---|---|---|---|---|---|---|
| ACE | 28 | 8140.52 | 1411 | | | | 8196.52 |
| AE | 20 | 8143.67 | 1419 | 3.15 | 8 | 0.925 | 8183.67 |
| CE | 20 | 8177.56 | 1419 | 37.04 | 8 | 0.000 | 8217.56 |
| E | 12 | 8271.47 | 1427 | 130.95 | 16 | 0.000 | 8295.47 |

Note: A = additive genetic, C = common or shared environment, E = non-shared environment, ep = number of estimated parameters, -2LL = -2 x log-likelihood, Δ-2LL = change in -2 x log-likelihood, Δdf = change in degrees of freedom, AIC = Akaike Information Criteria

Table S5. Pairwise polyserial phenotypic correlations for the predicted brain age difference (PBAD) scores.

| | 1. | 2. | 3. | 4. |
|---|---|---|---|---|
| 1. PBAD 51 to 55 | 1 | | | |
| 2. PBAD 56 to 60 | 0.69 | 1 | | |
| 3. PBAD 61 to 65 | 0.77 | 0.74 | 1 | |
| 4. PBAD 66 to 72 | 0.65 | 0.71 | 0.75 | 1 |

Table S6. Predicted brain age difference (PBAD) univariate model fitting results along with standardized variance components and 95% confidence intervals components for the best-fitting additive genetic & non-shared environment univariate ‘AE’ models.

| PBAD 51 to 55 | ep | -2LL | df | Δ-2LL | Δdf | p | AIC | A | C | E |
|---|---|---|---|---|---|---|---|---|---|---|
| ACE | 4 | 1409.56 | 224 | | | | 1417.56 | | | |
| AE | 3 | 1411.00 | 225 | 1.44 | 1 | 0.230 | 1417.00 | 0.64 (0.55-0.71) | - | 0.36 (0.29- 0.45) |
| CE | 3 | 1416.86 | 225 | 7.30 | 1 | 0.007 | 1422.86 | | | |
| E | 2 | 1509.11 | 226 | 99.56 | 2 | <0.001 | 1513.11 | | | |
| PBAD 56 to 60 | ep | -2LL | df | Δ-2LL | Δdf | p | AIC | A | C | E |
| ACE | 4 | 2593.15 | 440 | | | | 2601.15 | | | |
| AE | 3 | 2593.43 | 441 | 0.28 | 1 | 0.599 | 2599.43 | 0.71 (0.61-0.79) | - | 0.29 (0.21- 0.39) |
| CE | 3 | 2613.05 | 441 | 19.90 | 1 | <0.001 | 2619.05 | | | |
| E | 2 | 2669.24 | 442 | 76.09 | 2 | <0.001 | 2673.24 | | | |
| PBAD 61 to 65 | ep | -2LL | df | Δ-2LL | Δdf | p | AIC | A | C | E |
| ACE | 4 | 2569.07 | 413 | | | | 2577.07 | | | |
| AE | 3 | 2570.27 | 414 | 1.20 | 1 | 0.274 | 2576.27 | 0.59 (0.46-0.69) | - | 0.41 (0.31- 0.54) |
| CE | 3 | 2583.11 | 414 | 14.04 | 1 | <0.001 | 2589.11 | | | |
| E | 2 | 2624.11 | 415 | 55.04 | 2 | <0.001 | 2628.11 | | | |
| PBAD 66 to 72 | ep | -2LL | df | Δ-2LL | Δdf | p | AIC | A | C | E |
| ACE | 4 | 2096.56 | 346 | | | | 2104.56 | | | |
| AE | 3 | 2096.77 | 347 | 0.20 | 1 | 0.652 | 2102.77 | 0.60 (0.46-0.70) | - | 0.40 (0.30- 0.54) |
| CE | 3 | 2102.33 | 347 | 5.77 | 1 | 0.016 | 2108.33 | | | |
| E | 2 | 2143.32 | 348 | 46.76 | 2 | 0.000 | 2147.32 | | | |

Note: A = additive genetic, C = common or shared environment, E = non-shared environment, ep = number of estimated parameters, -2LL = -2 x log-likelihood, Δ-2LL = change in -2 x log-likelihood, Δdf = change in degrees of freedom, AIC = Akaike Information Criteria

Table S7. Predicted brain age difference (PBAD) multivariate model fitting results & comparisons between the saturated 'ACE' correlated factors model (reference), autoregression, 1- & 2-factor common pathways, & the independent pathways sub-models.

| | ep | -2LL | df | AIC | Δ-2LL | Δdf | p |
|---|---|---|---|---|---|---|---|
| ACE correlated factors (reference) | 34 | 8112.44 | 1405 | 8180.44 | | | |
| Autoregression | 18 | 8141.56 | 1421 | 8177.56 | 29.12 | 16 | 0.023 |
| 2-factor common pathway | 32 | 8118.33 | 1409 | 8182.33 | 5.89 | 4 | 0.208 |
| 1-factor common pathway | 23 | 8129.63 | 1417 | 8175.63 | 17.19 | 12 | 0.143 |
| Independent pathways | 28 | 8133.56 | 1411 | 8189.56 | 21.12 | 6 | 0.002 |

Note: ep = number of estimated parameters, -2LL = -2 x log-likelihood, Δ-2LL = change in -2 x log-likelihood, Δdf = change in degrees of freedom, AIC = Akaike Information Criteria

Table S8. Predicted brain age difference (PBAD) multivariate model fitting results & comparisons between the best fitting ACE 1-factor common pathway model and the AE, CE & E sub-models.

| | ep | -2LL | df | AIC | Δ-2LL | Δdf | p |
|---|---|---|---|---|---|---|---|
| ACE | 23 | 8129.63 | 1417 | 8175.63 | | | |
| AE | 18 | 8135.21 | 1422 | 8171.21 | 5.58 | 5 | 0.349 |
| CE | 18 | 8171.27 | 1422 | 8207.27 | 41.64 | 5 | <0.001 |
| E | 13 | 8304.07 | 1427 | 8330.07 | 174.43 | 10 | <0.001 |

Note: A = additive genetic, C = common or shared environment, E = non-shared environment, ep = number of estimated parameters, -2LL = -2 x log-likelihood, Δ-2LL = change in -2 x log-likelihood, Δdf = change in degrees of freedom, AIC = Akaike Information Criteria

Table S9. Predicted brain age difference (PBAD) additive genetic (below diagonal) & non-shared environmental latent factor correlations based on the best fitting AE 1-factor common pathway model.

| | 1. | 2. | 3. | 4. |
|---|---|---|---|---|
| 1. PBAD 51 to 55 | 1 | 0.49 | 0.46 | 0.47 |
| 2. PBAD 56 to 60 | 0.81 | 1 | 0.55 | 0.57 |
| 3. PBAD 61 to 65 | 0.93 | 0.87 | 1 | 0.54 |
| 4. PBAD 66 to 72 | 0.82 | 0.78 | 0.89 | 1 |

OpenMx R code for multivariate analysis of predicted brain aging (PBA)

```
# ACE correlated factors (reference) model
# Autoregression
# 2-factor common pathway
# 1-factor common pathway
# Independent pathways
# Multivariate model comparisons

# ACE correlated factors (reference) model
selVars   <- c("PBA_51_T1","PBA_56_T1","PBA_61_T1","PBA_66_T1",
                "PBA_51_T2","PBA_56_T2","PBA_61_T2","PBA_66_T2")
mzdata          <- subset(df2, zyg2019==1, selVars)
dzdata          <- subset(df2, zyg2019==2, selVars)
polycor::hetcor(dzdata, use="pairwise.complete.obs")
nv        = length(selVars)/2
ntv       = nv*2
thVals    = 0.5
aLabs     = paste("a",rev(nv+1-sequence(1:nv)),rep(1:nv,nv:1),sep="")
cLabs     = paste("c",rev(nv+1-sequence(1:nv)),rep(1:nv,nv:1),sep="")
eLabs     = paste("e",rev(nv+1-sequence(1:nv)),rep(1:nv,nv:1),sep="")
avals           = matrix(c( 20, 0, 0, 0,
                           0,25, 0, 0,
                           0, 0,20, 0,
                           0, 0, 0,20),nv, byrow=T)
cvals           = matrix(c( 5,0,0,0,
                           0,5,0,0,
                           0,0,5,0,
                           0,0,0,4),nv, byrow=T)
evals           = matrix(c( 7,0,0,0,
                           0,12,0,0,
                           0,0,10,0,
                           0,0,0,10),nv, byrow=T)
frees           = matrix(c( T,T,T,T,
                           T,T,T,T,
                           T,T,T,T,
                           T,T,T,T),nv, byrow=T)
multi_PBA = mxModel("ACE",
mxModel("top",
mxMatrix(       name="Means",     type="Full", nrow=1, ncol=nv, free=T, labels=c("m1","m2","m3","m4"),values = 0),
mxAlgebra(      name="expMean", cbind(Means,Means)),
mxMatrix(       name="A",               type="Symm", nrow=nv, ncol=nv, free=frees, values=avals, labels=aLabs, lbound= -5, ubound=40),
mxMatrix(       name="C",               type="Symm", nrow=nv, ncol=nv, free=frees, values=cvals, labels=cLabs, lbound=-20, ubound=40),
mxMatrix(       name="E",               type="Symm", nrow=nv, ncol=nv, free=frees, values=evals, labels=eLabs, lbound= -5, ubound=40),
mxAlgebra(      name="expCovMZ",expression= rbind( cbind(A+C+E, A+C  ), cbind(  A+C, A+C+E))),
mxAlgebra(      name="expCovDZ",expression= rbind( cbind(A+C+E, 0.5%x%A+C), cbind( 0.5%x%A+C, A+C+E) )),
mxAlgebra(      name="corrP",     expression=cov2cor(A+C+E)),
mxAlgebra(      name="corrA",     expression=cov2cor(A)),
mxAlgebra(      name="corrE",     expression=cov2cor(E)),
mxCI(c("corrA[2,1]","corrA[3,1]","corrA[3,2]","corrA[4,2]","corrA[4,3]", "corrE[2,1]","corrE[3,1]","corrE[3,2]","corrE[4,2]","corrE[4,3]",
```

```
        "VC[1,13]","VC[2,14]","VC[3,15]","VC[4,16]","VC[1,21]","VC[2,22]","VC[3,23]","VC[4,24]")),
 mxAlgebra(        name="VC",                  expression=cbind(A,C,E,A/(A+C+E),C/(A+C+E),E/(A+C+E)), dimnames=list(rep('VC',nv),rep(c('A','C','E','SA','SC','SE'),each=nv))) ),
 mxModel("MZ", mxData(             mzdata, type ="raw"), mxExpectationNormal("top.expCovMZ", means="top.expMean", dimnames=selVars), mxFitFunctionML() ),
 mxModel("DZ", mxData(             dzdata, type ="raw"), mxExpectationNormal("top.expCovDZ", means="top.expMean", dimnames=selVars), mxFitFunctionML() ),
 mxFitFunctionMultigroup(c("MZ","DZ")))
 omxGetParameters(multi_PBA)

# ACE
 summary( multi_PBA_fit       <- mxTryHard( multi_PBA,extraTries=25, greenOK=FALSE,checkHess=FALSE,intervals=F) )
 summary( multi_PBA_fit )
# AE model
 multi_PBA_AE                          <- mxModel( multi_PBA,name="AE")
 multi_PBA_AE                          <- omxSetParameters( multi_PBA_AE, cLabs, free=F, values=0)
 multi_PBA_AE                          <- omxSetParameters( multi_PBA_AE, c("a11","a22","a33","a44"), free=T, values=20)
 summary( multi_PBA_AE_fit  <- mxTryHard(            multi_PBA_AE, extraTries=25, greenOK=TRUE,checkHess=FALSE,intervals=F))
 summary( multi_PBA_AE_fit )

# CE model
 multi_PBA_CE                          <- mxModel( multi_PBA,name="CE")
 multi_PBA_CE                          <- omxSetParameters( multi_PBA_CE, aLabs, free=F, values=0)
 summary( multi_PBA_CE_fit  <- mxTryHard(            multi_PBA_CE, extraTries=5, greenOK=TRUE,checkHess=FALSE,intervals=T))
 summary( multi_PBA_CE_fit )
# E model
 multi_PBA_E                           <- mxModel( multi_PBA,name="E")
 multi_PBA_E                           <- omxSetParameters( multi_PBA_E, c(aLabs,cLabs),free=F,values=0)
 multi_PBA_E                           <- omxSetParameters( multi_PBA_E, labels=eLabs, free=T,lbound=0, ubound=40)
 summary( multi_PBA_E_fit    <- mxTryHard(            multi_PBA_E, extraTries=25, greenOK=TRUE,checkHess=FALSE,intervals=T))
 summary( multi_PBA_E_fit )
# Comparisons
 subs                                        <- c( multi_PBA_AE_fit, multi_PBA_CE_fit, multi_PBA_E_fit )
 comps                                       <- mxCompare( multi_PBA_fit,subs );comps
```

**# Autoregression**

```
selVars   <- c("PBA_51_T1","PBA_56_T1","PBA_61_T1","PBA_66_T1",
                 "PBA_51_T2","PBA_56_T2","PBA_61_T2","PBA_66_T2")
 mzdata          <- subset(df2, zyg2019==1, selVars)
 dzdata          <- subset(df2, zyg2019==2, selVars)
 cov(mzdata[selVars],use="complete")
 cov(dzdata[selVars],use="complete")
 describe(mzdata[selVars])
 describe(dzdata[selVars])
 nv        = length(selVars)/2
 ntv       = nv*2
 nVariables       = nv
 nFactors = nv
 psi_lab_a        = c("ia11","ia22","ia33","ia44")
 psi_lab_c        = c("ic11","ic22","ic33","ic44")
 psi_lab_e        = c("ie11","ie22","ie33","ie44")
 res_lab_a        = c("res_a1","res_a2","res_a3","res_a4")
 res_lab_c        = c("res_c1","res_c2","res_c3","res_c4")
```

```
res_lab_e              = "res_e"
betaF      = matrix(c(F,F,F,F,
                          T,F,F,F,
                          F,T,F,F,
                          F,F,T,F),nv,byrow = TRUE)
b_lab                  = matrix(c(NA, NA, NA, NA,
                         "b", NA, NA, NA,
                          NA,"b", NA, NA,
                          NA, NA,"b", NA),nv,byrow = TRUE)
b_labC                 = matrix(c(NA, NA, NA, NA,
                         "bC", NA, NA, NA,
                          NA,"bC", NA, NA,
                          NA, NA,"bC", NA),nv,byrow = TRUE)
b_labE                 = matrix(c(NA, NA, NA, NA,
                         "bE", NA, NA, NA,
                          NA,"bE", NA, NA,
                          NA, NA,"bE", NA),nv,byrow = TRUE)
loadS                  = diag(nFactors)
loadF                  = F
psi_a_val              = c( 21,3,5,5)
psi_c_val= c( -1,-1,-1,-1)
psi_e_val              = c( 3,4,-1,-1)
epsi_e_val             = 0.8

auto_PBA = mxModel("ACE",
 mxModel("top",
 mxMatrix( name="Mean",          type="Full", nrow=nv, ncol=1, free = T, labels = c("m1","m2","m3","m4"), values=0, lbound = -5, ubound = 5 ),
 mxAlgebra(name="expMean",               expression= cbind(t(lamba %*% (solve(I-beta) %*% (Mean))), t(lamba %*% (solve(I-beta) %*% (Mean)))) ),
 mxMatrix(name="psi_a",          type="Diag", nrow = nFactors, ncol = nFactors, free = T, labels = psi_lab_a, values = psi_a_val, lbound = -5,  ubound = 50 ),
 mxMatrix(name="psi_c",          type="Diag", nrow = nFactors, ncol = nFactors, free = T, labels = psi_lab_c, values = psi_c_val, lbound = -20, ubound = 20 ),
 mxMatrix(name="psi_e",          type="Diag", nrow = nFactors, ncol = nFactors, free = T, labels = psi_lab_e, values = psi_e_val, lbound = -20, ubound = 20 ),
 mxMatrix(name="beta",                   type="Full", nrow = nFactors, ncol = nFactors, free = betaF, labels = b_lab, lbound = -2.5, ubound = 2.5 ),
 mxMatrix(name="I",                              type="Iden", nrow = nFactors, ncol = nFactors),
 mxMatrix(name="lamba",                  type="Full", nrow = nv, ncol = nv, free = F, values = diag(nv)),
 mxMatrix(name="epsilon_e",  type="Diag", nrow = nVariables, ncol = nVariables, free = T, labels = res_lab_e, values = 4),
 mxAlgebra(name="A",                     expression= lamba %&% (solve(I-beta) %&% psi_a) ),                       # + epsilon_a
 mxAlgebra(name="C",                     expression= lamba %&% (solve(I-beta) %&% psi_c) ),                       # + epsilon_c
 mxAlgebra(name="E",                     expression= lamba %&% (solve(I-beta) %&% psi_e) + epsilon_e),        #
 mxAlgebra(name="expCovMZ",      expression= rbind( cbind(A+C+E, A+C), cbind(A+C, A+C+E))),
 mxAlgebra(name="expCovDZ",      expression= rbind( cbind(A+C+E, 0.5%x%A+C), cbind(0.5%x%A+C, A+C+E))),
 mxAlgebra(name="corrP",         expression= cov2cor(A+C+E)),
 mxAlgebra(name="corrA",         expression= cov2cor(A)),
 mxAlgebra(name="corrE",         expression= cov2cor(E)),
 mxAlgebra(name="VC",                    expression=cbind(A,C,E,A/(A+C+E),C/(A+C+E),E/(A+C+E)), dimnames=list(rep('VC',nv),rep(c('A','C','E','SA','SC','SE'),each=nv))),
 mxCI(c("ia11","ia22","ia33","ia44","ic11","ic22","ic33","ic44","ie11","ie22","ie33","ie44","res_e", "b") )),
 mxModel("MZ", mxData(                   mzdata, type ="raw"), mxExpectationNormal("top.expCovMZ", means="top.expMean", dimnames=selVars ), mxFitFunctionML() ),
 mxModel("DZ", mxData(                   dzdata, type ="raw"), mxExpectationNormal("top.expCovDZ", means="top.expMean", dimnames=selVars ), mxFitFunctionML() ),
 mxFitFunctionMultigroup(c("MZ","DZ")))
 omxGetParameters(auto_PBA)
 mxCheckIdentification( auto_PBA )
```

```
# ACE
 summary( auto_PBA_fit                    <- mxTryHard( auto_PBA,  extraTries=35, greenOK=FALSE,checkHess=FALSE,intervals=F) )
 summary( auto_PBA_fit )
# AE model
 auto_PBA_AE                                             <- mxModel( auto_PBA_fit,name="AE")
 auto_PBA_AE                                             <- omxSetParameters( auto_PBA_AE, labels=psi_lab_c, free=F, values=0)
 auto_PBA_AE                                             <- omxAssignFirstParameters( auto_PBA_AE )
 summary( auto_PBA_AE_fit                 <- mxTryHard(                    auto_PBA_AE, extraTries=25, greenOK=TRUE,checkHess=FALSE,intervals=F))
 summary( auto_PBA_AE_fit )
# CE model
 auto_PBA_CE                                             <- mxModel( auto_PBA_fit,name="CE")
 auto_PBA_CE                                             <- omxSetParameters( auto_PBA_CE, labels=c(psi_lab_a), free=F, values=0)
 auto_PBA_CE                                             <- omxSetParameters( auto_PBA_CE, labels=c(psi_lab_c), free=T, values=10)
 summary( auto_PBA_CE_fit                 <- mxTryHard(                    auto_PBA_CE, extraTries=15, greenOK=TRUE,checkHess=FALSE,intervals=F))
# E model
 auto_PBA_E                                              <- mxModel( auto_PBA_fit,name="E")
 auto_PBA_E                                              <- omxSetParameters( auto_PBA_E, labels=c(psi_lab_a,psi_lab_c), free=F, values=0)
 summary( auto_PBA_E_fit                  <- mxTryHard(                    auto_PBA_E, extraTries=15, greenOK=TRUE,checkHess=FALSE))
# Comparisons
 subs                                                           <- c(auto_PBA_AE_fit, auto_PBA_CE_fit, auto_PBA_E_fit )
 comps                                                          <- mxCompare( auto_PBA_fit,subs );comps

# 2-factor common pathway
selVars <- c("PBA_51_T1","PBA_56_T1","PBA_61_T1","PBA_66_T1",
             "PBA_51_T2","PBA_56_T2","PBA_61_T2","PBA_66_T2")
 mzdata          <- subset(df2, zyg2019==1, selVars)
 dzdata          <- subset(df2, zyg2019==2, selVars)
 nv      = length(selVars)/2
 ntv     = nv*2
 nVariables      = 3
 nFactors = 2
 psi_lab_a       = c("ia11","ia21","ia22")
 psi_lab_c       = c("ic11","ic21","ic22")
 psi_lab_e       = c("ie11","ie21","ie22")
 psi_a_val       = 0.7
 psi_c_val= 0.2
 psi_e_val       = 0.2
 res_lab_a       = c("res_a1","res_a2","res_a3","res_a4")
 res_lab_c       = c("res_c1","res_c2","res_c3","res_c4")
 res_lab_e       = c("res_e1","res_e2","res_e3","res_e4")
 loadS           = 0.8
 loadF           = T
 lamba_lab       = c("f11","f21","f31","f41", "f12","f22","f32","f42")

cp_2F_PBA = mxModel("ACE",
 mxModel("top",
  mxMatrix(name="Mean",              type="Full", nrow=1, ncol=nv, free = T, labels = c("m1","m2","m3","m4"), lbound = -10, ubound = 5 ),
  mxAlgebra(name="expMean",          expression= cbind(Mean, Mean)),
  mxMatrix(name="psi_a",             type="Symm", nrow = nFactors, ncol = nFactors, free = T, labels = psi_lab_a, values = 0, lbound = -40, ubound = 40), #
```

```
  mxMatrix(name="psi_c",                         type="Symm", nrow = nFactors, ncol = nFactors, free = T, labels = psi_lab_c, values = 0, lbound = -40, ubound = 40), #
  mxMatrix(name="psi_e",                         type="Symm", nrow = nFactors, ncol = nFactors, free = T, labels = psi_lab_e, values = 1, lbound = -40, ubound = 40), #
  mxMatrix(name="lamba",                         type="Full", nrow = nv, ncol = nFactors, free = c(T,T,T,T,F,T,T,T), labels= lamba_lab, values = 1, lbound = 0, ubound = 20), #
  mxMatrix(name="epsilon_a",                     type="Diag", nrow = nv, ncol = nv, free = T, labels = res_lab_a, values =   0, lbound = -40, ubound = 40), #
  mxMatrix(name="epsilon_c",                     type="Diag", nrow = nv, ncol = nv, free = T, labels = res_lab_c, values =   0, lbound = -40, ubound = 40), #
  mxMatrix(name="epsilon_e",                     type="Diag", nrow = nv, ncol = nv, free = T, labels = res_lab_e, values = 0.1, lbound = -40, ubound = 40), #
  mxAlgebra(name="A",                                    expression= lamba %&% psi_a + epsilon_a),
  mxAlgebra(name="C",                                    expression= lamba %&% psi_c + epsilon_c),
  mxAlgebra(name="E",                                    expression= lamba %&% psi_e + epsilon_e),
  mxAlgebra(name="expCovMZ",             expression= rbind( cbind(A+C+E, A+C), cbind(A+C, A+C+E))),
  mxAlgebra(name="expCovDZ",             expression= rbind( cbind(A+C+E, 0.5%x%A+C), cbind(0.5%x%A+C, A+C+E))),
  mxAlgebra(name="corrP",                expression= cov2cor(A+C+E)),
  mxAlgebra(name="corrA",                expression= cov2cor(A)),
  mxAlgebra(name="corrE",                expression= cov2cor(E)),
  mxMatrix(name="unitM",type="Unit", nrow=2, ncol=1),
  mxConstraint(name="ConVar",expression=diag2vec(psi_a+psi_c+psi_e)==unitM),
  mxCI(c("ia11","ia21","ia22","ie11","ie21","ie22",  "res_a1","res_a2","res_a3","res_a4","res_e1","res_e2","res_e3","res_e4", "f11","f21","f31","f41", "f12","f22","f32","f42")),
  mxAlgebra(name="VC",expression=cbind(A,C,E,A/(A+C+E),C/(A+C+E),E/(A+C+E)), dimnames=list(rep('VC',nv),rep(c('A','C','E','SA','SC','SE'),each=nv)))),
  mxModel("MZ", mxData(       mzdata, type ="raw"), mxExpectationNormal(         covariance="top.expCovMZ", means="top.expMean", dimnames=selVars ), mxFitFunctionML()),
  mxModel("DZ", mxData( dzdata, type ="raw"), mxExpectationNormal( covariance="top.expCovDZ", means="top.expMean", dimnames=selVars ), mxFitFunctionML()),
  mxFitFunctionMultigroup(c("MZ","DZ")))
  omxGetParameters(cp_2F_PBA)

# ACE
 summary( cp_2F_PBA_fit                  <- mxTryHardOrdinal(cp_2F_PBA,  extraTries=25, greenOK=FALSE,checkHess=FALSE,fit2beat=Inf,intervals=F) )
 summary( cp_2F_PBA_fit )
# AE model
 cp_2F_PBA_AE                                            <- mxModel( cp_2F_PBA_fit,name="AE")
 cp_2F_PBA_AE                                            <- omxSetParameters( cp_2F_PBA_AE, label=psi_lab_c,free=F,values= 0)
 cp_2F_PBA_AE                                            <- omxSetParameters( cp_2F_PBA_AE, label=res_lab_c,free=F,values= 0)
 cp_2F_PBA_AE                                            <- omxSetParameters( cp_2F_PBA_AE, label=psi_lab_a,free=T,values=10)
 cp_2F_PBA_AE                                            <- omxSetParameters( cp_2F_PBA_AE, label=res_lab_a,free=T,values=10)
 summary( cp_2F_PBA_AE_fit                   <- mxTryHard(                      cp_2F_PBA_AE, extraTries=25, greenOK=TRUE,checkHess=FALSE,intervals=F))
# CE model
 cp_2F_PBA_CE                                            <- mxModel( cp_2F_PBA_fit,name="CE")
 cp_2F_PBA_CE                                            <- omxSetParameters( cp_2F_PBA_CE, label=psi_lab_a,free=F,values= 0)
 cp_2F_PBA_CE                                            <- omxSetParameters( cp_2F_PBA_CE, label=res_lab_a,free=F,values= 0)
 cp_2F_PBA_CE                                            <- omxSetParameters( cp_2F_PBA_CE, label=psi_lab_c,free=T,values= 5)
 cp_2F_PBA_CE                                            <- omxSetParameters( cp_2F_PBA_CE, label=res_lab_c,free=T,values=10)
 summary( cp_2F_PBA_CE_fit                   <- mxTryHard(                      cp_2F_PBA_CE, extraTries=25, greenOK=TRUE,checkHess=FALSE,intervals=F))
# E model
 cp_2F_PBA_E                                             <- mxModel( cp_2F_PBA_fit,name="E")
 cp_2F_PBA_E                                             <- omxSetParameters( cp_2F_PBA_E, label=psi_lab_a,free=F,values= 0)
 cp_2F_PBA_E                                             <- omxSetParameters( cp_2F_PBA_E, label=psi_lab_c,free=F,values= 0)
 cp_2F_PBA_E                                             <- omxSetParameters( cp_2F_PBA_E, label=res_lab_a,free=F,values= 0)
 cp_2F_PBA_E                                             <- omxSetParameters( cp_2F_PBA_E, label=res_lab_c,free=F,values= 0)
# Compare models
  subs                                                           <- c( cp_2F_PBA_AE_fit,cp_2F_PBA_CE_fit, cp_2F_PBA_E_fit)
  comps                                                  <- mxCompare( cp_2F_PBA_fit,subs ); comps
```

**# 1-factor common pathway**

```
 selVars <- c("PBA_51_T1","PBA_56_T1","PBA_61_T1","PBA_66_T1",
                      "PBA_51_T2","PBA_56_T2","PBA_61_T2","PBA_66_T2")
 mzdata                  <- subset(df2, zyg2019==1, selVars); # dim(mzdata) # 475  10
 dzdata                  <- subset(df2, zyg2019==2, selVars); # dim(dzdata) # 335  10
 nv          = length(selVars)/2
 ntv         = nv*2
 nVariables              = 3
 nFactors = 1
 psi_lab_a               = c("ia11")
 psi_lab_c               = c("ic11")
 psi_lab_e               = c("ie11")
 psi_a_val               = 0.7
 psi_c_val= 0.2
 psi_e_val               = 0.2
 res_lab_a               = c("res_a1","res_a2","res_a3","res_a4")
 res_lab_c               = c("res_c1","res_c2","res_c3","res_c4")
 res_lab_e               = c("res_e1","res_e2","res_e3","res_e4")
 loadS                   = 0.8
 loadF                   = T
 lamba_lab               = c("f11","f21","f31","f41")

cp_1F_PBA = mxModel("ACE",
 mxModel("top",
  mxMatrix(name="Mean",                     type="Full", nrow=1, ncol=nv, free = T, labels = c("m1","m2","m3","m4"), lbound = -15, ubound = 5 ),
  mxAlgebra(name="expMean",                 expression= cbind(Mean, Mean)),
  mxMatrix(name="psi_a",                    type="Symm", nrow = nFactors, ncol = nFactors, free = T, labels = psi_lab_a, values = 0, lbound = -40, ubound = 40), #
  mxMatrix(name="psi_c",                    type="Symm", nrow = nFactors, ncol = nFactors, free = T, labels = psi_lab_c, values = 0, lbound = -40, ubound = 40), #
  mxMatrix(name="psi_e",                    type="Symm", nrow = nFactors, ncol = nFactors, free = T, labels = psi_lab_e, values = 1, lbound = -40, ubound = 40), #
  mxMatrix(name="lamba",                    type="Full", nrow = nv, ncol = nFactors, free = c(T,T,T,T), labels= lamba_lab, values = 1, lbound = 0, ubound = 20), #
  mxMatrix(name="epsilon_a",                type="Diag", nrow = nv, ncol = nv, free = T, labels = res_lab_a, values =   0, lbound = -40, ubound = 40), #
  mxMatrix(name="epsilon_c",                type="Diag", nrow = nv, ncol = nv, free = T, labels = res_lab_c, values =   0, lbound = -40, ubound = 40), #
  mxMatrix(name="epsilon_e",                type="Diag", nrow = nv, ncol = nv, free = T, labels = res_lab_e, values = 0.1, lbound = -40, ubound = 40), #
  mxAlgebra(name="A",                               expression= lamba %&% psi_a + epsilon_a),
  mxAlgebra(name="C",                               expression= lamba %&% psi_c + epsilon_c),
  mxAlgebra(name="E",                               expression= lamba %&% psi_e + epsilon_e),
  mxAlgebra(name="expCovMZ",                expression= rbind( cbind(A+C+E, A+C), cbind(A+C, A+C+E))),
  mxAlgebra(name="expCovDZ",                expression= rbind( cbind(A+C+E, 0.5%x%A+C), cbind(0.5%x%A+C, A+C+E))),
  mxAlgebra(name="corrP",                   expression= cov2cor(A+C+E)),
  mxAlgebra(name="corrA",                   expression= cov2cor(A)),
  mxAlgebra(name="corrE",                   expression= cov2cor(E)),
  mxMatrix(name="unitM",type="Unit", nrow=nFactors, ncol=nFactors),
  mxConstraint(name="ConVar",expression=diag2vec(psi_a+psi_c+psi_e)==unitM),
  mxCI(c("ia11","ie11", "res_a1","res_a2","res_a3","res_a4","res_e1","res_e2","res_e3","res_e4", "f11","f21","f31","f41",
  "VC[1,13]","VC[2,14]","VC[3,15]","VC[4,16]", "VC[1,21]","VC[2,22]","VC[3,23]","VC[4,24]","corrA[2,1]")),
  mxAlgebra(name="VC",expression=cbind(A,C,E,A/(A+C+E),C/(A+C+E),E/(A+C+E)), dimnames=list(rep('VC',nv),rep(c('A','C','E','SA','SC','SE'),each=nv)))),
  mxModel("MZ", mxData(       mzdata, type ="raw"), mxExpectationNormal(         covariance="top.expCovMZ", means="top.expMean", dimnames=selVars ), mxFitFunctionML()),
  mxModel("DZ", mxData( dzdata, type ="raw"), mxExpectationNormal( covariance="top.expCovDZ", means="top.expMean", dimnames=selVars ), mxFitFunctionML()),
  mxFitFunctionMultigroup(c("MZ","DZ")))
  omxGetParameters(cp_1F_PBA)
```

```
# ACE
 summary( cp_1F_PBA_fit                    <- mxTryHardOrdinal(cp_1F_PBA,  extraTries=25, greenOK=FALSE,checkHess=FALSE,fit2beat=Inf,intervals=F) )
# AE model
 cp_1F_PBA_AE                                        <- mxModel( cp_1F_PBA_fit,name="AE")
 cp_1F_PBA_AE                                        <- omxSetParameters( cp_1F_PBA_AE, label=psi_lab_c,free=F,values= 0)
 cp_1F_PBA_AE                                        <- omxSetParameters( cp_1F_PBA_AE, label=res_lab_c,free=F,values= 0)
 cp_1F_PBA_AE                                        <- omxSetParameters( cp_1F_PBA_AE, label=c("f11","f21","f31","f41"),              free=T,values=5, ubound=10)
 cp_1F_PBA_AE                                        <- omxSetParameters( cp_1F_PBA_AE, label=c(res_lab_a,res_lab_e),              free=T,values=0.5, ubound=20)
 cp_1F_PBA_AE                                        <- omxSetParameters( cp_1F_PBA_AE, label="ia11",free=T,values=20,ubound=40)
 summary( cp_1F_PBA_AE_fit                    <- mxTryHard(                     cp_1F_PBA_AE, extraTries=50, greenOK=TRUE,checkHess=FALSE,intervals=F))
# CE model
 cp_1F_PBA_CE                                        <- mxModel( cp_1F_PBA_fit,name="CE")
 cp_1F_PBA_CE                                        <- omxSetParameters( cp_1F_PBA_CE, label=psi_lab_a,free=F,values= 0)
 cp_1F_PBA_CE                                        <- omxSetParameters( cp_1F_PBA_CE, label=res_lab_a,free=F,values= 0)
 cp_1F_PBA_CE                                        <- omxSetParameters( cp_1F_PBA_CE, label=c("f11","f21","f31","f41"), free=T,values=5, ubound=10)
 cp_1F_PBA_CE                                        <- omxSetParameters( cp_1F_PBA_CE, label=c(res_lab_c,res_lab_e), free=T,values=0.5, ubound=20)
 cp_1F_PBA_CE                                        <- omxSetParameters( cp_1F_PBA_CE, label="ie11",free=T,values=20,ubound=40)
 summary( cp_1F_PBA_CE_fit                    <- mxTryHard(                     cp_1F_PBA_CE, extraTries=25, greenOK=TRUE,checkHess=FALSE,intervals=F))
# E model
 cp_1F_PBA_E                                         <- mxModel( cp_1F_PBA_fit,name="E")
 cp_1F_PBA_E                                         <- omxSetParameters( cp_1F_PBA_E, label=psi_lab_a,free=F,values= 0)
 cp_1F_PBA_E                                         <- omxSetParameters( cp_1F_PBA_E, label=psi_lab_c,free=F,values= 0)
 cp_1F_PBA_E                                         <- omxSetParameters( cp_1F_PBA_E, label=res_lab_a,free=F,values= 0)
 cp_1F_PBA_E                                         <- omxSetParameters( cp_1F_PBA_E, label=res_lab_c,free=F,values= 0)
 cp_1F_PBA_E                                         <- omxSetParameters( cp_1F_PBA_E, label=c("f11","f21","f31","f41"),  free=T,values=5, ubound=10)
 cp_1F_PBA_E                                         <- omxSetParameters( cp_1F_PBA_E, label="ie11",
      free=T,values=20,ubound=40)
 summary( cp_1F_PBA_E_fit               <- mxTryHard(                     cp_1F_PBA_E, extraTries=25, greenOK=TRUE,checkHess=FALSE,intervals=F))
# Compare models
  subs                                                     <- c( cp_1F_PBA_AE_fit,cp_1F_PBA_CE_fit,cp_1F_PBA_E_fit)
  comps                                              <- mxCompare( cp_1F_PBA_fit,subs ); comps

# Independent pathways
 selVars <- c("PBA_51_T1","PBA_56_T1","PBA_61_T1","PBA_66_T1",
                  "PBA_51_T2","PBA_56_T2","PBA_61_T2","PBA_66_T2")
 mzdata              <- subset(df2, zyg2019==1, selVars)
 dzdata              <- subset(df2, zyg2019==2, selVars)
 nv          = length(selVars)/2
 ntv         = nv*2
 nVariables          = 3
 nFactors = 1
 psi_lab_a           = c("ia11")
 psi_lab_c           = c("ic11")
 psi_lab_e           = c("ie11")
 psi_a_val           = 0.7
 psi_c_val= 0.2
 psi_e_val           = 0.2
 res_lab_a           = c("res_a1","res_a2","res_a3","res_a4")
 res_lab_c           = c("res_c1","res_c2","res_c3","res_c4")
```

```
 res_lab_e              = c("res_e1","res_e2","res_e3","res_e4")
 loadS                  = 0.8
 loadF                  = T
 lamba_lab_a            = c("f11_a","f21_a","f31_a","f41_a")
 lamba_lab_c            = c("f11_c","f21_c","f31_c","f41_c")
 lamba_lab_e            = c("f11_e","f21_e","f31_e","f41_e")
 FT        = matrix(c(T,F,F,F,
                      T,F,F,F,
                      T,F,F,F,
                      T,F,F,F),nv,byrow = TRUE)
 lamba_lab_a <- matrix(c("f11_a",NA,NA,NA,
                         "f21_a",NA,NA,NA,
                         "f31_a",NA,NA,NA,
                         "f41_a",NA,NA,NA),nv,byrow = TRUE)
 lamba_lab_c <- matrix(c("f11_c",NA,NA,NA,
                         "f21_c",NA,NA,NA,
                         "f31_c",NA,NA,NA,
                         "f41_c",NA,NA,NA),nv,byrow = TRUE)
  lamba_lab_e <- matrix(c("f11_e",NA,NA,NA,
                         "f21_e",NA,NA,NA,
                         "f31_e",NA,NA,NA,
                         "f41_e",NA,NA,NA),nv,byrow = TRUE)

ip_PBA = mxModel("ACE",
 mxModel("top",
  mxMatrix(name="Mean",            type="Full", nrow=1, ncol=nv, free = T, labels = c("m1","m2","m3","m4"), lbound = -15, ubound = 5 ),
  mxAlgebra(name="expMean",        expression= cbind(Mean, Mean)),
  mxMatrix(name="lamba_a",         type="Lower", nrow = nv, ncol = nv, free = FT, labels= lamba_lab_a, values = 1, lbound = -20, ubound = 20),
  mxMatrix(name="lamba_c",         type="Lower", nrow = nv, ncol = nv, free = FT, labels= lamba_lab_c, values = 1, lbound = -20, ubound = 20),
  mxMatrix(name="lamba_e",         type="Lower", nrow = nv, ncol = nv, free = FT, labels= lamba_lab_e, values = 1, lbound = -20, ubound = 20),
  mxMatrix(name="epsilon_a",       type="Diag", nrow = nv, ncol = nv, free = T, labels = res_lab_a, values =   0, lbound = -40, ubound = 40),
  mxMatrix(name="epsilon_c",       type="Diag", nrow = nv, ncol = nv, free = T, labels = res_lab_c, values =   0, lbound = -40, ubound = 40),
  mxMatrix(name="epsilon_e",       type="Diag", nrow = nv, ncol = nv, free = T, labels = res_lab_e, values = 0.1, lbound = -40, ubound = 40),
  mxAlgebra(name="A",                   expression= lamba_a %*% t(lamba_a) + epsilon_a %*% t(epsilon_a)),
  mxAlgebra(name="C",                   expression= lamba_c %*% t(lamba_c) + epsilon_c %*% t(epsilon_c)),
  mxAlgebra(name="E",                   expression= lamba_e %*% t(lamba_e) + epsilon_e %*% t(epsilon_e)),
  mxAlgebra(name="expCovMZ",       expression= rbind( cbind(A+C+E, A+C), cbind(A+C, A+C+E))),
  mxAlgebra(name="expCovDZ",       expression= rbind( cbind(A+C+E, 0.5%x%A+C), cbind(0.5%x%A+C, A+C+E))),
  mxAlgebra(name="VC",expression=cbind(A,C,E,A/(A+C+E),C/(A+C+E),E/(A+C+E)), dimnames=list(rep('VC',nv),rep(c('A','C','E','SA','SC','SE'),each=nv)))),
  mxModel("MZ", mxData(    mzdata, type ="raw"), mxExpectationNormal(      covariance="top.expCovMZ", means="top.expMean", dimnames=selVars ), mxFitFunctionML()),
  mxModel("DZ", mxData( dzdata, type ="raw"), mxExpectationNormal( covariance="top.expCovDZ", means="top.expMean", dimnames=selVars ), mxFitFunctionML()),
  mxFitFunctionMultigroup(c("MZ","DZ")))
  omxGetParameters(ip_PBA)
  mxCheckIdentification(ip_PBA)

# ACE
 summary( ip_PBA_fit               <- mxTryHardOrdinal(ip_PBA,  extraTries=25, greenOK=FALSE,checkHess=FALSE,fit2beat=Inf,intervals=F) )
# AE model
 ip_PBA_AE                         <- mxModel( ip_PBA_fit,name="AE")
 ip_PBA_AE                         <- omxSetParameters( ip_PBA_AE, label=c("f11_c","f21_c","f31_c","f41_c"),          free=F,values= 0)
```

```
 ip_PBA_AE                                     <- omxSetParameters( ip_PBA_AE, label=c("res_c1","res_c2","res_c3","res_c4"),free=F,values= 0)
 summary( ip_PBA_AE_fit          <- mxTryHard(              ip_PBA_AE, extraTries=50, greenOK=TRUE,checkHess=FALSE,intervals=F))
# CE model
 ip_PBA_CE                                     <- mxModel( ip_PBA_fit,name="CE")
 ip_PBA_CE                                     <- omxSetParameters( ip_PBA_CE, label=c("f11_a","f21_a","f31_a","f41_a"),           free=F,values= 0)
 ip_PBA_CE                                     <- omxSetParameters( ip_PBA_CE, label=c("res_a1","res_a2","res_a3","res_a4"),       free=F,values= 0)
 summary( ip_PBA_CE_fit          <- mxTryHard(              ip_PBA_CE, extraTries=25, greenOK=TRUE,checkHess=FALSE,intervals=F))
# E model
 ip_PBA_E                                      <- mxModel( ip_PBA_fit,name="E")
 ip_PBA_E                                      <- omxSetParameters( ip_PBA_E, c("f11_c","f21_c","f31_c","f41_c"),           free=F,values= 0)
 ip_PBA_E                                      <- omxSetParameters( ip_PBA_E, c("res_c1","res_c2","res_c3","res_c4"),       free=F,values= 0)
 ip_PBA_E                                      <- omxSetParameters( ip_PBA_E, c("f11_a","f21_a","f31_a","f41_a"),           free=F,values= 0)
 ip_PBA_E                                      <- omxSetParameters( ip_PBA_E, c("res_a1","res_a2","res_a3","res_a4"),       free=F,values= 0)
 summary( ip_PBA_E_fit           <- mxTryHard(              ip_PBA_E, extraTries=25, greenOK=TRUE,checkHess=FALSE,intervals=F))
# Compare models
   subs                                        <- c( ip_PBA_AE_fit,ip_PBA_CE_fit,ip_PBA_E_fit)
   comps                               <- mxCompare( ip_PBA_fit,subs ); comps

# Multivariate model comparisons
subs                                           <- c( auto_PBA_fit, cp_2F_PBA_fit, cp_1F_PBA_fit, ip_PBA_fit  )
comps                                          <- mxCompare( multi_PBA_fit, subs );comps
```
